# How cognitive genetic factors influence fertility outcomes: A mediational SEM analysis

**DOI:** 10.1101/070128

**Authors:** Michael A. Woodley Menie, Joseph A. Schwartz, Kevin M. Beaver

## Abstract

Utilizing a newly released cognitive Polygenic Score (PGS) from Wave IV of Add Health (*n* = 1,886), structural equation models (SEMs) examining the relationship between PGS and fertility (which is approximately 50% complete in the present sample), utilizing measures of verbal IQ and educational attainment as potential mediators, were estimated. The results of indirect pathway models revealed that verbal IQ mediates the positive relationship between PGS and educational attainment, and educational attainment in turn mediates the negative relationship between IQ and a latent fertility measure. The direct path from PGS to fertility was non-significant. The model was robust to controlling for age, sex and race, furthermore the results of a multi-group SEM revealed no significant differences in the estimated path coefficients across sex. These results indicate that those predisposed towards higher IQ by virtue of higher PGS values are also predisposed towards trading fertility against time spent in education, which contributes to those with higher PGS values producing fewer offspring.

---

Fertility in human populations is a complex phenomenon produced by a multifactorial arrangement of genetic and environmental antecedents (Tropf et al., 2015). Particular attention has been paid to the role of cognitive variables in determining fertility outcomes. Negative associations between cognitive ability and fertility were first intuited by Galton (1869), when he noted that individuals from lower socioeconomic status backgrounds tended to produce larger families than those from higher status backgrounds. Early studies utilizing direct measurements of cognitive ability (i.e. IQ tests) and sibling number (as an indicator of the reproductive success of the previous generation) corroborated Galton’s intuition, via the finding of negative correlations between the two (e.g. Lentz, 1927). These findings led to the prediction that IQ scores may decline over the course of generations due to selection pressure stemming from these negative associations. However, longitudinal studies comparing cohorts evaluated in the early decades of the 20^th^ century with those evaluated in subsequent decades found the opposite trend, i.e. an apparent increase in IQ (Cattell, 1950),consistent with what is now termed the Flynn effect.

Interest in the question of the causes and implications of the negative relationship between IQ and fertility subsequently waned, until the 1990s, when Lynn (1996) proposed that the selection pressure stemming from the negative association might be reducing the more heritable variance components of IQ (which he termed *genotypic IQ),* the effects of which are being masked by environmental improvements (such as increased access to micronutrients) enhancing the more environmentally sensitive variance components of IQ *(phenotypic IQ).* Lynn also suggested that educational attainment (essentially the amount of time spent in education) may be a major cause of this negative relationship, as higher ability individuals might trade fertility against the opportunity to acquire skills and knowledge *(somatic capital),* an effect that should be especially strong among women, given their relatively narrower fertility window. Consistent with this trade-off model, negative correlations have been directly observed between educational attainment and fertility, both across cultures (Meisenberg, 2008) and time (Skirbekk, 2008). Meisenberg (2010) utilized structural equations modelling to analyse factors mediating the relationship between IQ and fertility among a large, nationally representative sample of the US population. Consistent with the trade-off model, it was found that educational attainment partially mediated the negative relationship between the two. Meisenberg furthermore found indications of more strongly negative correlations between educational attainment and fertility among the female population.

Recent developments involving genome-wide association studies (GWAS) have yielded evidence that certain single nucleotide polymorphisms (SNPs; termed ‘hits’) have robust (genome-wide significant) associations with individual differences in educational attainment and also IQ (e.g. Reitveld et al., 2013, 2014). Increased power has led to the discovery of dozens of these ‘hits’, which when concatenated into polygenic ‘risk’ scores (PGS), seem to predict a portion of the individual variation in both educational attainment and IQ across several, large databases (Okbay et al.,2016; Selzam et al., 2016). One potential use for these PGS values is in investigating their role as antecedents of fertility. Two studies have attempted to do precisely this. Beauchamp (2016) utilized a large and representative sample of the population of the US born between 1931 and 1953 sourced from *The US Health and Retirement Study* (HRS). It was found using regression analysis that an educational attainment PGS negatively predicted fertility (measured as relative lifetime reproductive success). The magnitude of the association, coupled with a correction for the ‘missing’ heritability of educational attainment, led Beauchamp to predict that attained education should decline at a rate of 1.5 months per generation due to genetic selection. A second study (Conley et al. 2016) also utilizing data from the US HRS, but this time from samples born between 1920 and 1955 replicated Beauchamp’s finding of negative associations between an educational attainment PGS and completed fertility, however, the apparent secular increase in the strength of the negative phenotypic association between the two was not paralleled by a corresponding secular increase in the strength of the genetic association.

Unlike Meisenberg (2010), neither Beauchamp (2016) nor Conley and co-workers (2016) examined IQ directly, nor was the possibility of mediation considered. In point of fact, both Beauchamp (2016) and Conley and co-workers (2016) treat educational attainment as a phenotype, and as a potential target for selection. Educational attainment is better conceptualized however as an outcome of facultative calibration during childhood and young adulthood in response to the action of heritable characteristics such as IQ, with which it shares around 60 to 70 percent of its genetic variance (Okbay et al., 2016). Thus, as Lynn predicted, it is the trade-off between the opportunity to acquire somatic capital and fertility *evoked* by a high-IQ individual’s genetic affinity for the specific set of environments embodied in the educational system, rather than a genetic disposition towards lower fertility *per se,* that should result in reduced fertility among such individuals. Consistent with this is the relatively low additive heritability of fertility in modern populations (≈.30; Tropf et al., 2015), which indicates considerable potential plasticity in response to environmental mediators such as years of schooling. Furthermore genes for higher IQ do not seem to supress intrinsic fertility, as indicated by the presence of small-magnitude positive phenotypic correlations with indicators of reproductive fitness potential, such as sperm quality (Arden, Gottfredson, Miller & Pierce, 2009). This is also consistent with the observation that among Western population, in the absence of factors such as universal education prior to the 19^th^ century, potential proxies for IQ such as social status actually seem to have been positively phenotypically correlated with fitness outcomes (Skirbekk, 2008).

In the present study, the question of the potential mediating role of educational attainment on the relationship between IQ, PGS and fertility will be considered in relation to newly released cognitive PGS data from the National Longitudinal Study of Adolescent to Adult Health *(Add Health),* as described in a recent publication by Domingue and co-workers (2015).

## Methods

### Data

Data for the current study were drawn from the *Add Health,* a longitudinal and nationally representative sample of American youth enrolled in middle or high school during the 1994-1995 academic year (Harris et al., 2009). Participants were recruited using a multi-stage cluster design resulting in the selection of 132 schools (80 high schools and 52 middle schools) stratified by region, urbanicity, school type, ethnicity, and size. Students at each selected school were asked to participate in the study, with a total of over 90,000 students completing the in-school portion. A subsample of these students were asked to participate in the more comprehensive in-home portion of the study. In total, 20,744 participants aged between 12 and 21 (and 17,700 of their primary caregivers) completed the Wave I in-home interview, which covered a wide variety of topics including physical and mental health, delinquent behavior, physical development, and family interactions. The second wave of the study was completed between 1995 and 1996 and included 14,738 participants that also completed the Wave I interview. Wave III of the study was completed between 2001 and 2002, when participants were aged between 18 and 28 years old, and included 15,197 of the original participants. Based on the amount of time that elapsed between the Wave II and Wave III interviews (approximately 5-6 years) the survey instruments utilized were updated to tap more age-appropriate topics such as romantic relationships, civic participation, and contact with the criminal justice system. Finally, the fourth, and most recent wave of interviews, were completed between 2007 and 2008 when participants were between 24 and 34 years old. A total of 15,701 participants from the in-home sample completed the Wave IV interview.

Nested within the full sample of in-home participants is a subsample of over 3,000 pairs of individuals from the same household (including twin and sibling pairs; for more information on sampling procedures and the composition of the sibling subsample, see Harris et al., 2006). Members of the sibling subsample who participated in the fourth wave of the study were also asked to provide a saliva sample for DNA extraction.^1^ Approximately 96 percent of the sibling subsample provided saliva samples (for additional details, see Harris et al., 2013). After a series of quality control (QC) and quality assurance (QA) steps, genome-wide data were made available for a total of 1,886 participants (905 males and 981 females; for a more detailed description of the QC/QA procedures see McQueen et al., 2015). The final analytic sample was restricted to those individuals with valid genome-wide information (n = 1,886). Compared to omitted participants, members of the final analytic sample had a significantly greater number of children (*t* = -5.27, *p* < .001), pregnancies (or a significant other who had been pregnant; *t* = -5.00, *p* < .001), and number of live births (*t* = -5.29, *p* < .001), but had significantly lower levels of verbal IQ (t = 4.11, *p* < .001) and educational attainment (t = 6.29, *p* < .001), and were younger (*t* = 3.41, *p* < .001). The final analytic sample also included a significantly greater proportion of Caucasian participants (χ^2^ = 55.46, *p* < .001), but the proportion of male participants was not significantly different from those Wave IV participants that were excluded from the final analytic sample (χ^2^ = 1.85, *p* = .17). Means,proportions, minimum and maximum values, and sample sizes for all measures included in the current study are presented in Table 1. Descriptive statistics are presented for the full sample as well as sex-restricted subsamples.

### Measures

#### Fertility

Fertility was measured using three items from the Wave IV interview tapping reproductive success. More specifically, female participants were asked to report the number of times they had been pregnant (including a current pregnancy and pregnancies that resulted in an abortion, miscarriage, or stillbirth), the total number of live births that resulted from such pregnancies, and the total number of living children at the time of the interview. Male participants were asked a similar set of questions, but the pregnancy item was modified to reflect the number of times each respondent had “made a partner pregnant.” Female participants reported a greater number of living children (*M* = 1.38, SD = 1.25; *t* = 6.69, *p* < .001), pregnancies (*M* = 1.83, SD = 1.66; *t* = 5.86, *p* < .001), and live births (*M* = 1.28, SD = 1.26; *t* = 6.83, *p* < .001) compared to male participants (M = .98, SD = 1.20; *M* = 1.37, SD = 1.78; *M* = .89,SD = 1.19, respectively). The construction of the fertility measure is discussed in more detail below.

#### Verbal IQ

*Verbal IQ* was measured using the Picture Vocabulary Test (PVT), a modified version of the Peabody Picture Vocabulary Test-Revised (PPVT-R), which was administered during Wave I interviews. Scores on the PVT were measured continuously and standardized by age (*M* = 97.13, SD = 14.43). The difference in PVT scores between male (*M* = 97.88, SD = 14.57) and female (*M* = 96.44, SD = 14.27) participants was significant (*t* = -2.12, *p* = .03). Strictly speaking the PPVT test series measures verbal ability, however it has been found to correlate strongly with measures tapping other cognitive domains, indicating that it is a good measure of *general intelligence (g)* also (Dunn & Dunn, 1997).

#### Educational Attainment

During Wave IV interviews, participants were asked, “What is the highest level of education you have achieved to date?” Responses were coded categorically as follows: 1 = 8^th^ grade or less; 2 = some high school; 3 = high school graduate; 4 = some vocational school after high school; 5 = completed vocational school; 6 = some college; 7 = completed a four-year college degree; and 8 = additional college beyond a four-year degree (*M* = 5.45, SD = 1.85). Females reported significantly greater levels of educational attainment relative to males (*t* = 5.11, *p* < .001).

#### Polygenic Education Score

During Wave IV interviews, saliva was collected using the Oragene collection method and genotyping was conducted with the Illumina HumanOmni1-Quad v1 platform (Harris et al., 2012; McQueen et al., 2015). After QC procedures, valid data were available for 940,862 SNPs (for more information, see McQueen et al., 2014). Using information from the genome-wide database, polygenic education scores (PGS) were calculated for each participant. More specifically, SNPs included in the genetic database were matched with SNPs identified in a large-scale GWAS of educational attainment (Rietveld et al., 2013). Each of the identified SNPs were then multiplied by the effect size estimated in the GWAS, summed, and then z-transformed (*M* = 0, SD = 1) to create the PGS (for additional details, see Domingue et al., 2015).

#### Statistical Covariates

Three demographic covariates were also included in the estimated models. First, Wave 1 age was measured continuously in years (*M* = 16.02, SD = 1.72) and was not significantly different between males and females (*t* = -1.43, *p* = .15). Second, race was measured dichotomously such that 0 = Caucasian (54.24%) and 1 = all other races (45.76%). The distribution of the examined race categories was not significantly different across sex (χ^2^ = .22, *p* = .64). Controlling for race allowed for the potentially confounding effects of population stratification on the distribution of PGS and the outcome measures to be controlled. Third, sex was included in all models analyzing the full sample and coded 0 = female (52.01%) and 1 = male (47.99%).

#### Plan of Analysis

Models were estimated in three steps. First, zero-order correlations were estimated in an effort to examine the potential bivariate associations. The second step of the analysis involved a series of measurement models aimed at identifying a fertility factor. Confirmatory factor analysis (CFA) was used to identify the latent fertility measure using the three indicators from Wave IV (total number of pregnancies, live births, and children). Since the CFA included only three indicators, the resulting model was just-identified (i.e., the number of estimated parameters was equal to the degrees of freedom), resulting in uninterpretable fit parameters (e.g., (χ^2^ = .00, *p* = 1.00). Despite this limitation, the magnitude (and accompanying p-values) of the standardized factor loadings can be used as a secondary indicator of the extent to which the estimated model taps the underlying latent construct of fertility. Since the results of preliminary analyses revealed that all of the indicators included in the fertility measurement model significantly differed between males and females, a series of multi-group CFA (MGCFA) models were estimated to test for measurement invariance across sex. Comparative model fit was estimated using likelihood ratio tests (LRTs), the results of which are distributed in χ^2^ units.

The third and final step of the analysis involved the estimation of a series of structural equation models (SEMs) examining the direct and indirect pathways between the PGS, educational attainment, verbal IQ, and fertility. Based on the primary objectives of the current study, four specific pathways were examined. First, the direct pathway between the PGS and fertility was examined. Second, in an effort to identify the potential mechanisms that ultimately connect previously observed associations between PGS and fertility, two indirect pathways were examined, with verbal IQ and highest level of educational attainment included as potential mediators. An additional indirect pathway, in which educational attainment mediated the association between verbal IQ and fertility, was also examined. Third, in an effort to identify potential mechanisms that ultimately connect PGS and educational attainment, verbal IQ was also entered into the SEM as a potential intervening variable between the PGS and highest level of educational attainment. All models included controls for age, race, and sex (in models examining the full analytic sample). In an effort to examine potential moderating effects of sex, additional multi-group SEMSs were also estimated. The resulting indirect effects for males and females were compared using difference tests and z-scores. Directly in line with previous research (MacKinnon et al., 2002; 2004; Preacher & Hayes, 2008), the standard errors, and corresponding confidence intervals, for the indirect pathway models were estimated using bootstrapping procedures with 5,000 replications.

All models were estimated using *Mplus* 7.4 (Muthén & Muthén, 1998-2012). Measurement models were estimated using a maximum likelihood estimator with robust standard errors (MLR), while SEMs (due to the presence of a categorical educational attainment measure) were estimated using a weighted least squares estimator with robust standard errors (WLSMV) and missing data were handled using full information maximum likelihood (FIML) estimation. Model fit was assessed using χ^2^, the comparative fit index (CFI), the Tucker-Lewis Index (TLI), and the root mean square error of approximation (RMSEA). The following conventional cut-off values were used to evaluate acceptable model fit: CFI≥.95; TLI≥.95; and RMSEA ≤ .06 (for more information, see Hu & Bentler, 1999).

## Results

### Bivariate Correlations

The results of the zero-order correlations are presented as a heat map in Figure 1, along with the corresponding coefficients and accompanying significance levels. In an effort to aid in interpretation, darker colors indicate larger correlation coefficients. As indicated in the figure, all of the examined study variables were significantly (p < .001) associated with one another at the bivariate level. The fertility measures were all strongly associated with one another with correlations ranging between *r* = .76 (the association between pregnancies and live births) and *r* = .99 (the association between number of children and live births). In addition, the fertility measures were negatively associated with verbal IQ, educational attainment, and the PGS, providing preliminary evidence that the association between genetic influences and fertility may be mediated by educational influences. Finally, verbal IQ, educational attainment and the PGS were all modestly associated with one another.

### Measurement Models

The next step in the analysis involved the estimation of a series of measurement models assessing the latent fertility measure. The first measurement model included the full analytic sample and allowed the three fertility indicators to freely load on a single latent factor. As mentioned above, due to the limited number of indicator variables, this model is just-identified and yielded perfect (yet trivial) fit indices. Despite this limitation, a number of findings from the analyses provide tentative evidence that the indicator variables are tapping a single underlying latent construct. First, the results of the zero-order correlations indicated that the fertility measures were all strongly correlated with one another. Second, the standardized factor loadings from the measurement model were large, ranging between .760 (total number of pregnancies) and .997 (total number of live births), and statistically significant (*p* < .001). Third, a supplemental measurement model (results not presented, but available upon request) in which the resulting latent fertility factor was regressed on the statistical covariates (age, race, and sex) was estimated in an effort to increase overall degrees of freedom, allowing for the estimation of fit indices. Despite a significant χ^2^ estimate (which is sensitive to sample size), the resulting model provided an adequate fit to the data (χ^2^ = 26.58, df = 6, *p* < .001; CFI = .98; TLI =.95; RMSEA=.04).

Based on these findings, a series of MGCFA models were estimated to test for measurement invariance across sex. Subsequently restrictive models were tested against the configural model (i.e. all parameters freed across both groups) using LRTs. The resulting models indicated both metric (Δχ^2^ = 4.47, df = 2, *p* = .11) and scalar invariance (Δχ^2^ = 6.76, df = 4, *p* = .15). However, constraining the residual variances to equality (i.e., residual variance invariance) significantly worsened overall fit (Δχ^2^ = 14.00, df = 7, *p* = .05). Finally, constraining the variance of the fertility factor to equality across groups (i.e., factor variance invariance) did not worsen fit (Δχ^2^> = 8.55, df = 5, *p* = .13), but constraining factor means to equality resulted in worsened fit (Δχ^2^ = 47.74, df = 6, *p* < .001). Based on these results, the residual variances of the indicators and the means of the fertility factor were freely estimated across sex, but all other parameters were constrained to equality. The resulting MG-CFA fit the data closely (χ^2^ = 8.55, df = 5, *p* = .13; CFI = .99; TLI = .99; RMSEA = .03).

### Structural Equation Models

The next step in the analysis involved the estimation of a series of SEMs aimed at examining the direct and indirect pathways between the study measures. The SEM examining the full analytic sample, along with standardized path coefficients and accompanying robust standard errors, is presented in Figure 2. Since only theoretically relevant paths were estimated, the resulting model was over-identified, effectively allowing the estimation of fit indices (df = 11), with the resulting model providing an adequate fit to the data (χ^2^ = 28.23, df = 11, *p* < .01; CFI = .99; TLI = .99; RMSEA = .03). The results indicated a negative, but nonsignificant association between the PGS and fertility (*β* = -.045, *p* = .11), but positive and significant associations between the PGS and both verbal IQ (*β* = .19, *p* < .001) and educational attainment (*β* = .11, *p* < .001). Educational attainment was also significantly associated with fertility (*β* = -.28, *p* < .001), indicating that participants with higher levels of education had overall lower rates of fertility. The association between verbal IQ and fertility was also negative, but only marginally significant (β = -.05, *p* = .049). Finally, the association between verbal IQ and educational attainment was positive and significant (β = .39, *p* < .001).

The next step of the analysis involved the estimation of a multiple group model in an effort to examine direct and indirect pathways among the male and female subsamples. The multiple group model fit the data closely (χ^2^ = 30.28, df = 22, *p* = .11; CFI = .99; TLI = .99; RMSEA = .02) and the results are presented in Figure 3. The top panel of the figure presents the results for the male subsample. The overall pattern of results converged with findings from the full sample. The association between the PGS and fertility remained nonsignificant (*β* = -.07, *p* = .07), but the associations between the PGS and both verbal IQ (*β* = .21, *p* < .001) and educational attainment (*β* = .11, *p* = .004) were both positive and significant. In addition, the association between educational attainment and fertility was negative and significant (*β* = -.24, *p* < .001) and the association between verbal IQ and educational attainment was positive and significant (*β* = .40, *p* < .001). The primary divergence from the pattern of findings observed in the full sample was a nonsignificant association between verbal IQ and fertility (*β* = -.05, *p* = .27). The results from the female subsample are presented in the bottom panel of the figure and were highly similar to the findings from the male subsample. Once again, the association between the PGS and fertility was nonsignificant (*β* = -.03, *p* = .51), but the associations between the PGS and both verbal IQ (*β* = .18, *p* < .001) and educational attainment (*β* = .11, *p* < .001) were significant. Furthermore, the associations between educational attainment and both fertility (*β* = -.32, *p* < .001) and verbal IQ (*β* = .40, *p* < .001) were significant. Similar to the results from the male subsample, the association between verbal IQ and fertility was nonsignificant (*β* = -.06, *p* = .07).

The final step of the analysis involved the estimation of a series of indirect pathways aimed at examining the potential mechanisms that mediate the association between PGS and both educational attainment and fertility. The results of the indirect pathway models are presented in Table 2. The model examining the full sample provided an adequate fit to the data (χ^2^ = 28.23, df = 11, *p* = .003; CFI = .99; TLI = .99; RMSEA = .03) and revealed that verbal IQ significantly mediated the association between the PGS and educational attainment (*b* = .083, *p* < .001). Verbal IQ (*b* = -.011, *p* < .001) and educational attainment (*b* = -.033, *p* < .001) also significantly mediated the association between PGS and fertility. Finally, educational attainment also significantly mediated the association between verbal IQ and fertility (*b* = -.008, *p* < .001). The multiple group model also provided an adequate fit to the data (χ^2^ = 37.39, df = 25, *p* = .053; CFI = .99; TLI = .99; RMSEA = .02). The results for the male subsample revealed that verbal IQ significantly mediated the association between PGS and educational attainment (*b* = .088, *p* < .001). In addition, educational attainment significantly mediated the association between PGS and fertility *(b* = - .026, *p* = .002) and verbal IQ and fertility (*b* = -.006, *p* < .001). The association between PGS and fertility was not mediated by verbal IQ (*b* = -.010, *p* = .31). The results from the female subsample were similar, with verbal IQ significantly mediating the association between PGS and educational attainment (*b* = .079, *p* < .001), but did not significantly mediate the association between PGS and fertility (b= -.012, *p* = .12). Finally, educational attainment significantly mediated the association between PGS and fertility (*b* = -.039, *p* = .002) and the association between verbal IQ and fertility (*b* = -010, *p* < .001). The difference between the observed indirect effects for the male and female subsamples were compared using a z-score and are presented in the final columns of the table. The results of the difference tests revealed nonsignificant differences between all of the examined indirect effects, indicating that the examined pathways were not significantly moderated by sex.

## Discussion

The results of the SEMs were consistent with the expectation that verbal IQ should mediate the relationship between PGS and educational attainment. The direct effect of IQ on educational attainment is furthermore consistent with the expectation that the latter is (in part) an outcome variable of the former, rather than the other way around. This inference of causation accords with the finding that the direct effect of education on IQ is at the level of specific skills and competencies,rather than at the level of *g* (Ritchie et al., 2015), which is the more genetically substantive variance component of IQ (Rimfeld et al., 2015). Hence the path PGS → IQ → Education Attainment would appear to be justified in as much as the PVT test functions in these models as a good proxy for *g* (e.g. Dunn & Dunn, 1997). The presence of direct paths from PGS to both verbal IQ and educational attainment are furthermore consistent with the existence of substantial shared genetic variance between the two measures (Okbay et al., 2016).

Also consistent with expectations is the finding that educational attainment mediates the relationship between verbal IQ and the latent fertility measure. Importantly, the PGS did not significantly directly predict fertility once the pattern of mediation was taken into consideration. In this sample, the genetic disposition towards a given level of IQ does not therefore entail a direct fitness penalty. Instead, the negative correlation with fertility is largely a function of individuals with different cognitive genotypes actively seeking out different degrees of exposure to education, which is in turn traded against fertility. The significant direct path from verbal IQ to fertility (which was only found in the mixed-sex model) furthermore suggests the independent action of additional unmodelled IQ-dependent factors on fertility.

The results of the SEMs were robust to controlling for age, race and sex (in the first model). The results of a multiple group SEM furthermore revealed no significant indications of dimorphism in the strength of the path coefficients, which runs contrary to the expectation that females should be more sensitive to the effects of education on fertility.

One difference between the present study and the studies of Beauchamp (2016) and Conley and co-workers (2016) concerns the nature of the PGS variable. In the case of the present study, the PGS variable derives from the study of Rietveld et al. (2013), who found three loci to be reliable predictors of educational attainment. The PGS variable employed by Beauchamp in particular was sourced from a more recent and much larger GWAS analysis (Okbay et al., 2016), which found associations between educational attainment and 74 loci – a considerably larger number. Examining the pattern of mediation using the more recent PGS would be an important step in determining the robustness of these results.

Another important difference between the present study and those of Beauchamp and Conley and co-workers concerned their use of a sample that was in completed fertility (the participants of the subsample of the *Health and Retirement Study* utilized by Beauchamp for example were aged between 50 and 70 years). The mean age of our Wave IV participants by contrast is 28.4 years (with ages ranging from 24 to 34 years). Fertility is typically complete by the mid-40’s for both males and females in the West (although males can continue to produce children into their sixth decade of life, in practice relatively few of them do so, with only around 2.5% of males continuing or starting to produce children after the age of 45; Boschini et al., 2011). At age 29, males in the US have completed approximately 51.5% and females 50.8% of their fertility (see: Martinez et al., 2012, Tables 3 and 4, pp. 15-16). This is important as it has been noted that incomplete fertility inflates the negative association between fertility and IQ, largely because those with higher IQ are typically older when they start producing offspring, thus a larger proportion of high-IQ individuals will register as childless in their 20’s than in their 40’s (Neiss et al., 2002; Vining, 1982, 1995.) It is thus possible that the present results may have been somewhat different had the sample been in completed fertility. The effect of education as a mediator might weaken, as those with higher-IQs (and who have spent greater amounts of time in education) will have had an opportunity to reproduce, whereas other factors, such as income, might also come to play a significant and independent role in determining fertility outcomes at completed fertility (Meisenberg, 2010). It is also expected that the sex difference would be more prominent at completed fertility as the larger variance in male reproductive years enables those with higher-IQ to continue reproducing for longer - often with younger and also lower-IQ female partners who have relatively higher fertility, which has the effect of attenuating the negative IQ-fertility association among males to a greater degree than among females (Meisenberg, 2010). We nevertheless expect the basic pattern of mediation detected in the present study to persist at completed fertility. This expectation is consistent with Meisenberg’s (2010) finding of a role for educational attainment as a mediator of the phenotypic IQ-fertility relationship in a representative sample of the US population much closer to having completed its fertility (aged between 39 and 47 years).

This study is the first to take advantage of newly available data on cognitive genetics in attempting to examine the various causal and meditational pathways linking IQ to fertility that have been proposed in previous phenotype-only research. Future efforts should therefore focus on replicating the present findings using other databases in which PGS data along with measures of IQ, educational attainment and completed fertility are available.

## Footnotes

1 All Wave IV participants (*n* = 15,701) were asked to provide saliva samples, but the polygenic education scores employed in the current study (more detail provided below) were only estimated for the sibling subsample (for more information see Domingue et al., 2015; Harris et al., 2013; McQueen et al., 2015).

## Reference

Arden, R., Gottfredson, L.S., Miller, G., & Pierce, A. (2009). Intelligence and semen quality are positively correlated. Intelligence, 37, 277–282.

Beauchamp, J.P. (2016). Genetic evidence for natural selection in humans in the contemporary United States. Proceedings of the National Academy of Sciences USA, 113, 7774–7779.

Boschini, A., Håkanson, C., Rosén, Å., & Sjögren, A. (2011). Trading off or having it all? Completed fertility and mid-career earnings of Swedish men and women. Uppsala: IFAU—Institute for labour market policy evaluation. Working Paper.

Cattell, R.B. (1950). The fate of national intelligence: test of a thirteen-year prediction. The Eugenics Review, 42, 136–148.

Conley, D., Laidley, T., Belsky, D.W., Fletcher, J.M., Boardman, J.D., & Domingue B.W. (2016). Assortative mating and differential fertility by phenotype and genotype across the 20^th^ century. Proceedings of the National Academy of Sciences USA, 113, 6647–6652.

Domingue, B.W., Belsky, D.W., Conley, D., Harris, K.M., Boardman, J.D. (2015). Polygenic influence on educational attainment: New evidence from the National Longitudinal Study of Adolescent Health. AERA Open, 1, 1.13.

Dunn, L.M., & Dunn, L.M. (1997). Peabody Picture Vocabulary Test-III. Circle Pines, MD: American Guidance Service.

Galton, F. (1869). Hereditary genius. London: MacMillan.

Harris, K.M., Florey, T., Tabor, J., Bearman, P.S., Jones, J., & Udry, J.R. (2009). The national longitudinal study of adolescent health: Research design. Available from http://www.cpc.unc.edu/projects/addhealth/design.

Harris, K.M., Halpern, C.T., Haberstick, B.C., & Smolen, A. (2013). The National Longitudinal Study of Adolescent Health (Add Health) sibling pairs data. Twin Research and Human Genetics, 16, 391–398.

Harris, K. M., Halpern, C. T., Smolen, A., & Haberstick, B. C. (2006). The National Longitudinal Study of Adolescent Health (Add Health) twin data. Twin Research and Human Genetics, 9, 988–997.

Hu, L.T., & Bentler, P.M. (1999). Cutoff criteria for fit indexes in covariance structure analysis: Conventional criteria versus new alternatives. Structural Equation Modeling: A Multidisciplinary Journal, 6, 1–55.

Lentz, T. (1927). Relation of IQ to size of family. Journal of Educational Psychology, 18, 486–496.

Lynn, R. (1996). Dysgenics: Genetic deterioration in modern populations. Westport: Praeger.

MacKinnon, D.P., Lockwood, C.M., Hoffman, J.M., West, S.G., & Sheets, V. (2002). A comparison of methods to test mediation and other intervening variable effects. Psychological Methods, 7, 83–104.

MacKinnon, D.P., Lockwood, C.M., & Williams, J. (2004). Confidence limits for the indirect effect: Distribution of the product and resampling methods. Multivariate Behavioral Research, 39, 99–128.

Martinez, G., Daniels, K., & Chandra, A. (2012). Fertility of men and women aged 15-44 years in the United States: National survey of family growth, 2006-2010. National Health Statistics Report, 51, 1–28.

McQueen, M.B., Boardman, J.D., Domingue, B.W., Smolen, A., Tabor, J., Killeya-Jones, L., • & Harris, K.M. (2015). The national longitudinal study of adolescent to adult health (add health) sibling pairs genome-wide data. Behavior Genetics, 45, 12–23.

Meisenberg, G. (2010). The reproduction of intelligence. Intelligence, 38, 220–230.

Meisenberg, G. (2008). How universal is the negative correlation between education and fertility? Journal of Social, Political and Economic Studies, 33, 205–227.

Muthén, L. K., & Muthén, B. O. (2008–2012). Mplus user’s guide (7th ed.). Los Angeles, CA: Muthén & Muthén

Neiss, M., Rowe, D.C., & Rodgers, J.L. (2002). Does education mediate the relationship between IQ and age of first birth? A behavioural genetic analysis.Journal of Biosocial Science, 34, 259–276.

Okbay, A., Beauchamp, J.P., Fontana, M.A., Lee, J.J., Tune, H.P., Rietveld, C.A• Benjamin, D.J. (2016). Genome-wide association study identifies 74 loci associated with educational attainment. Nature. Doi:10.1038/nature17671

Preacher, K.J., & Hayes, A.F. (2008). Asymptotic and resampling strategies for assessing and comparing indirect effects in multiple mediator models. Behavior Research Methods, 40, 879–891.

Rimfeld, K., Kovas, Y., Dale, P.S., & Plomin, R. (2015). Pleiotropy across academic subjects at the end of compulsory education. Scientific Reports, 5:11713.

Rietveld, C.A., Medland, S.E., Derringer, J., Yang, J., Esko, T., Martin, N.W• Koellinger, P.D. (2013). GWAS of 126,559 individuals identifies genetic variants associated with educational attainment. Science, 340, 1467–1471.

Rietveld, C.A., Esko, T., Davies, G., Pers, T.H., Turley, P., Benyamin, B• Koellinger, P.D. (2014). Common genetic variants associated with cognitive performance identified using the proxy-phenotype method. Proceedings of the National Academy of Sciences, USA, 111, 13790–13794.

Ritchie, S.J., Bates, T.C., & Deary, I.J. (2015). Is education associated with improvements in general cognitive ability, or in specific skills? Developmental Psychology, 51, 573–582.

Selzam, S., Krapohl, E., von Stumm, S., O’Reilly, P.F., Rimfeld, K., Kovas, Y• Plomin, R. (2016). Predicting educational achievement from DNA. Molecular Psychaitry. Doi: 10.1038/mp.2016.107.

Skirbekk, V. (2008). Fertility trends by social status. Demographic Research, 18, 145–180.

Tropf, F.C., Stulp, G., Barban, N., Visscher, P.M., Yang, J., Sneider, H., & Mills, M.C. (2015). Human fertility, molecular genetics, and natural selection in modern societies. PLOS ONE, 10, e0126821.

Vining, D.R. (1982). On the possibility of the re-emergence of a dysgenic trend with respect to intelligence in American fertility differentials. Intelligence, 6, 241–264.

Vining, D.R. (1995). On the possibility of the re-emergence of a dysgenic trend with respect to intelligence in American fertility differentials: an update. Personality and Individual Differences, 19, 259–263.

